# Molecular mechanisms of uric acid transport by the native human URAT1 and its inhibition by anti-gout drugs

**DOI:** 10.1101/2024.09.11.612394

**Authors:** Canrong Wu, Chao Zhang, Sanshan Jin, James Jiqi Wang, Antao Dai, Jiuyin Xu, Heng Zhang, Xuemei Yang, Xinheng He, Qingning Yuan, Wen Hu, Youwei Xu, Mingwei Wang, Yi Jiang, Dehua Yang, H. Eric Xu

## Abstract

Gout, a common and painful disease, stems from hyperuricemia, where elevated blood uric acid levels lead to urate crystal formation in joints and kidneys. The human urate transporter 1 (hURAT1) plays a critical role in urate homeostasis by facilitating urate reabsorption in the renal proximal tubule, making it a key target for gout therapy. Pharmacological inhibition of hURAT1 with drugs such as dotinurad, benzbromarone, lesinurad, and verinurad promotes uric acid excretion and alleviates gout symptoms. Here we present cryo-electron microscopy structures of native hURAT1 bound with these anti-gout drugs in the inward-open state, and with uric acid in inward-open, outward-open, and occluded states. Complemented by mutagenesis and cell-based assays, these structures reveal the mechanisms of uric acid reabsorption and hURAT1 inhibition. Our findings elucidate the molecular basis of uric acid transport and anti-gout medication action, and provide a structural framework for the rational design of next-generation therapies for hyperuricemia and gout.

## Introduction

Gout, a prevalent and debilitating form of inflammatory arthritis, manifests through the deposition of monosodium urate crystals in joints and tissues^1^. This condition primarily results from hyperuricemia, defined as an excessive accumulation of uric acid in the bloodstream. With the global incidence of both gout and hyperuricemia escalating, these conditions pose a growing public health challenge. In the United States, the prevalence of gout has risen dramatically over the past few decades, affecting approximately 3.9% of the adult population, or about 9.2 million people^2^. Additionally, hyperuricemia affects about 20% of general populations^3^. This condition significantly contributes to the development of gout and related health complications. The increasing prevalence, coupled with the significant impact on quality of life and healthcare costs, underscores the urgent need for improved therapeutic strategies.

Initially asymptomatic, sustained hyperuricemia can evolve into gout, promote the formation of kidney stones, and lead to renal failure^4^. The therapeutic landscape for hyperuricemia involves two primary strategies: inhibition of uric acid production and enhancement of uric acid excretion. Xanthine oxidase inhibitors (XOIs), such as allopurinol and febuxostat, function by reducing uric acid synthesis^5^. However, they do not address the excretion issues that are problematic in approximately 90% of hyperuricemia cases, where patients suffer from inadequate uric acid clearance^6^. Additionally, XOIs are associated with severe side effects, including hypersensitivity reactions, hepatotoxicity, and renal impairment, particularly in patients with pre-existing renal conditions or those intolerant to XOIs^7^.

The human urate transporter 1 (URAT1) is a crucial member of the SLC22A subfamily, which includes both organic anion transporters (OATs) and organic cation transporters (OCTs)^8^. Encoded by the SLC22A12 gene, URAT1 plays an essential role in uric acid homeostasis^9^. It is primarily expressed in the apical membrane of the proximal tubule epithelial cells in the kidney, where it functions as a high-affinity urate-anion exchanger. This transporter facilitates the reabsorption of uric acid from the glomerular filtrate back into the bloodstream, a process critical for maintaining serum uric acid levels within a physiological range.

Genetic studies have underscored the importance of URAT1 in uric acid regulation. Loss-of-function mutations in SLC22A12 lead to renal hypouricemia, a condition characterized by excessive urinary uric acid excretion and very low serum uric acid levels^10,11^. Conversely, certain polymorphisms in this gene have been associated with hyperuricemia and increased risk of gout, highlighting the critical role of URAT1 in urate homeostasis^12^.

Given its pivotal function in urate reabsorption, URAT1 has emerged as a key target for pharmacological intervention in the treatment of hyperuricemia and gout. URAT1 inhibitors offer a direct method to enhance uric acid excretion by blocking its reabsorption in the kidneys. Early URAT1 inhibitors such as benzbromarone and lesinurad, however, were limited by their low selectivity and activity, leading to adverse reactions and restricted clinical use in some regions^13–15^. Recent advancements have led to the development of more selective inhibitors. Verinurad, for example, is currently undergoing promising Phase III clinical trials, demonstrating high selectivity and efficacy^16^. Additionally, dotinurad has been approved in several countries, showing good tolerability and effectiveness^17,18^.

Despite significant strides in developing URAT1 inhibitors, the instability and low expression levels of the human URAT1 has prevented the structure determination, and the lack of structural information has impeded a deeper understanding of drug action mechanisms and selectivity. In this study, we overcame the technical difficulties in expression and purification of the native human URAT1 and used cryo-electron microscopy (cryo-EM) to unveil the structures of URAT1 bound with four anti-gout drugs (dotinurad, benzbromarone, lesinurad, and verinurad), revealing their binding and inhibition mechanisms. Additionally, we resolved native URAT1 in three conformations with uric acid, shedding light on the substrate transport mechanism. These insights provide a structural basis for designing next-generation URAT1 inhibitors, enhancing both the efficacy and selectivity of treatments for hyperuricemia and gout.

## Results

### The overall structure of URAT1

To elucidate the structural basis of URAT1 function and inhibition, we expressed full-length wild-type human URAT1 in HEK 293E cells and purified it for cryo-EM analysis (Figure S1). Because of the instability and low expression levels of the human URAT1, we had to express it in large scales, which only yielded less than 0.01 mg per liter of HEK 293E cells. In addition, we had screened an extensive set of various detergents for cryo-EM studies that ultimately led to an optimal detergent-lipid combination of LMNG, GDN, and cholesterol (see methods). These efforts allowed us to complex URAT1 with its natural substrate, uric acid, and four structurally diverse anti-gout drugs: lesinurad, verinurad, benzbromarone, and dotinurad.

We successfully resolved cryo-EM structures of hURAT1 bound to the four anti-gout drugs in the inward-open state at resolutions ranging from 3.2 to 3.6 Å (Figure 1, A to C; Figure S2 to S5; and table S1). Additionally, we determined structures of URAT1 in complex with uric acid in three distinct conformational states: inward-open (3.3 Å), outward-open (4.1 Å), and occluded (4.7 Å) (Figures 1C, S2 to S5, and Table S1).

**Figure 1.**
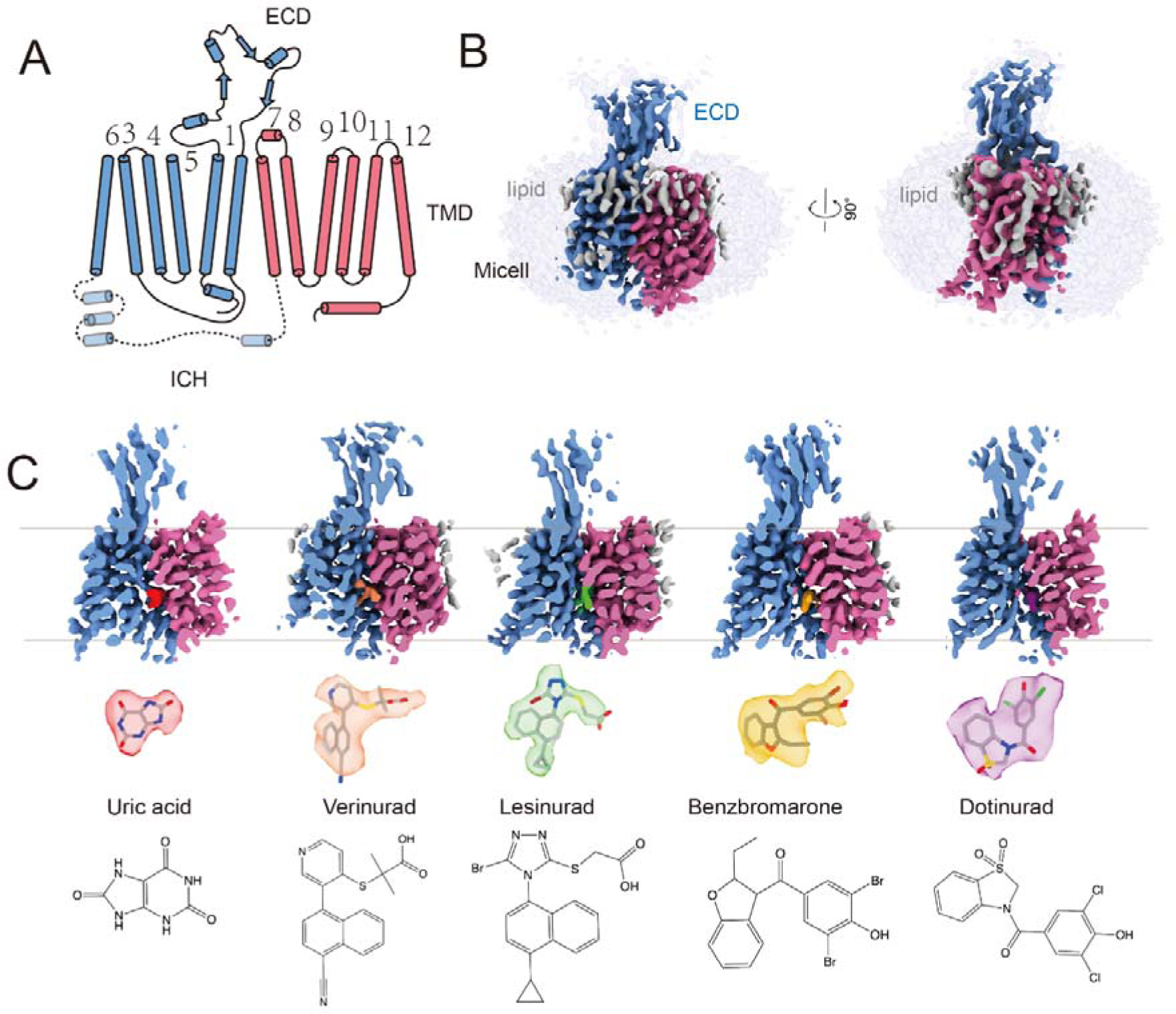
Cryo-EM structures of URAT1. **(A).** Predicted diagram of the structural elements of URAT1, highlighting three distinct regions: the transmembrane domain (TMD), the extracellular domain (ECD), and a predicted intracellular helical (ICH) bundle. The pseudo two-fold symmetric repeats, TM1-6 (blue) and TM7-12 (red), are depicted. **(B).** Orthogonal views of the density map for URAT1 (blue and red) in the inward-open state within the micelle (gray, 90% transparency). Robust lipid densities surrounding the TMD of URAT1 are marked in gray. **(C).** Cross-sectional view of the density map for uric acid-, verinurad-, lesinurad-, benzbromarone-, and dotinurad-bound URAT1 (from left to right) in the inward-open state. The colors of the substrate and inhibitors are indicated, with their densities and structural formulas highlighted correspondingly.

The five inward-open structures exhibited remarkably similar conformations, with root mean square deviation (RMSD) values for Cα atoms between 0.11 to 0.35 Å. The overall architecture of URAT1 comprises three distinct regions: the transmembrane domain (TMD), the extracellular domain (ECD), and a predicted intracellular helical (ICH) bundle, which was not resolved in our structures (Figure 1A and 1B).

The TMD of URAT1 displays a characteristic Major Facilitator Superfamily (MFS) topology, consisting of 12 transmembrane helices arranged in two pseudo-symmetric halves: the N-terminal domain (NTD, TM1–TM6) and the C-terminal domain (CTD, TM7–TM12). The inward-facing conformation of the URAT1 TMD bears resemblance to that of rOAT1 and OCT1^19–21^, two related transporters from the SLC22A family, with RMSD values of 1.14 and 1.57 Å, respectively (Figure 2A). However, URAT1 exhibits a more compact structure compared to rOAT1, with a smaller distance between the cytoplasmic ends of the NTD and CTD. Notably, helices TM4, TM5, TM8, and TM10 in URAT1 are displaced inward by 2.3-5.6 Å relative to their positions in rOAT1 (Figure 2A).

**Figure 2.**
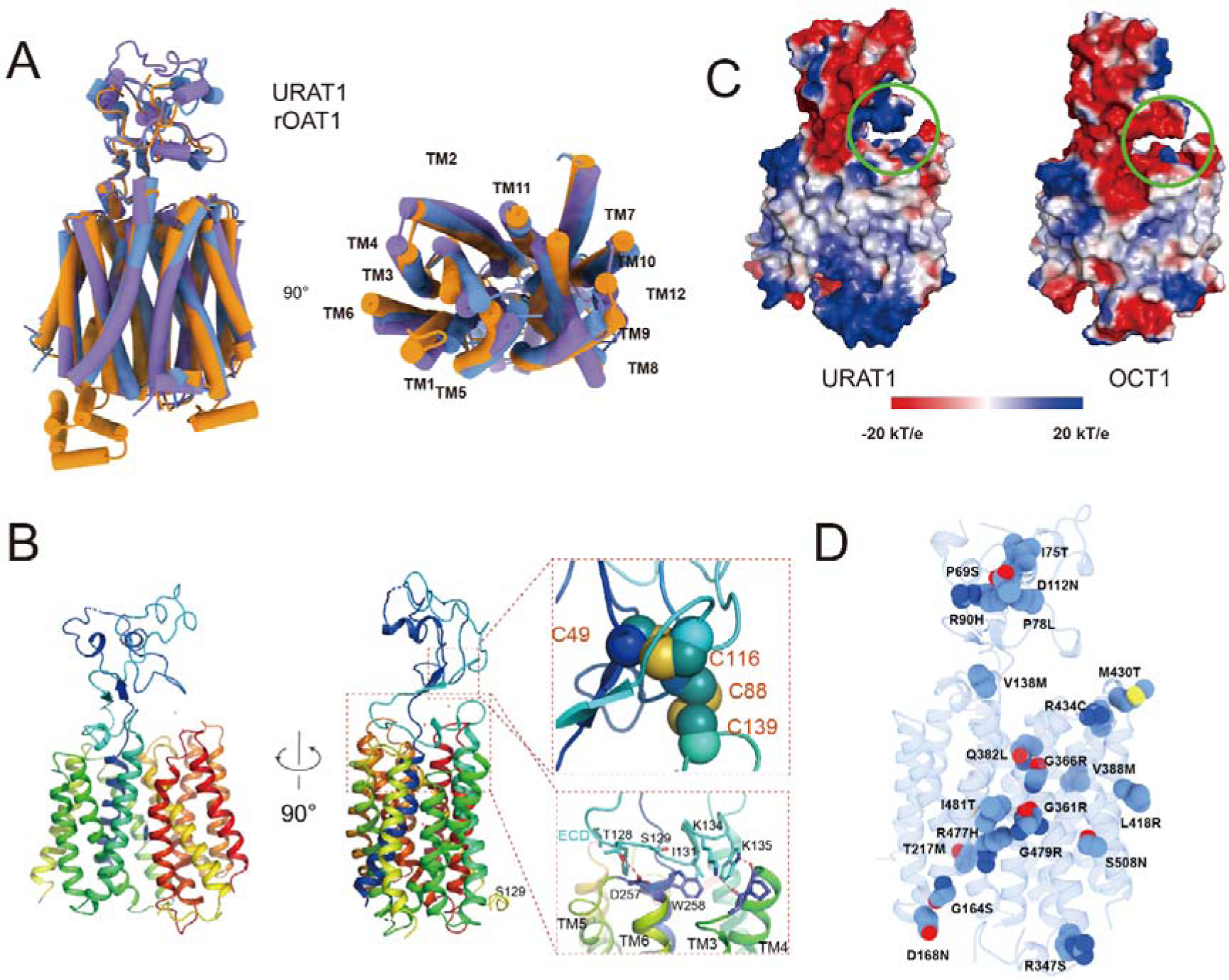
Overall structure of URAT1. (A) Structural comparisons of the inward-facing conformation of URAT1 with rOAT1 (PDB: 8SDY) and OCT1 (PDB: 8SC1). The N-terminal domain (NTD) was used as a reference for structural alignments. (B) Cartoon representation of URAT1 highlighting the ECD and the ECD-TMD interface. Residues involved in ECD-TMD interactions are displayed, and cysteines participating in disulfide bond formation are shown as spheres. (C) Electrostatic potential surfaces of URAT1 and OCT1 calculated in PyMOL (red to blue, −20LJkT/e to +20LJkT/e). The extracellular domain (ECD) and extracellular portions of the transmembrane domain (TMD) bundle in URAT1 and OCT1 are shown. The circled region is positively charged in URAT1 and negatively charged in OCT1. (D) Mapping of natural URAT1 variants onto the structure of URAT1. Residues associated with natural human URAT1 variants are displayed as spheres.

A distinctive feature of the SLC22A family is the extended ECD located between TM1 and TM2. The ECD’s structural integrity is maintained by two conserved disulfide bonds:

C49-C116, which links two antiparallel beta-sheets, and C88-C139, which connects the ECD to the apex of TM2 (Figure 2B). These disulfide bonds are a common feature among SLC22 family members. In URAT1, the ECD can be only observed in inward open conformation (Figure S4), at this state, the ECD forms a cap-like structure over the N-lobe, stabilized by loops between TM3–TM4 and TM5–TM6 (Figure 2B).

Interestingly, the ECD and the extracellular portions of the TMD bundle in URAT1 form a positively charged cavity (circle area in Figure 2C). This feature may play a crucial role in capturing organic anions such as uric acid, contrasting with the predominantly negatively charged cavity in OCT1, which is specialized for binding organic cations (Figure 2C). This structural characteristic provides insight into the substrate specificity of URAT1 and its function in uric acid transport.

URAT1 exhibits a high degree of genetic polymorphism, with natural variants widely distributed across the transmembrane domain (TMD), extracellular domain (ECD), and the unresolved intracellular domain (ICD) (Figure 2D). Mutations in these regions can significantly affect substrate recognition, conformational transitions, or protein expression levels of URAT1, potentially leading to conditions such as renal hypouricemia ^10^.

### Recognition of uric acid by URAT1

Having determined the overall structure of URAT1, we next focused on elucidating the molecular basis of its substrate recognition. The 3.3 Å resolution structure of URAT1 in complex with uric acid revealed critical details about the substrate binding site. The density maps clearly showed uric acid bound within a pocket formed by residues from transmembrane helices TM5, TM7, TM8, and TM10 (Figure 3A). This binding pocket is characterized by an aromatic cage composed of phenylalanine residues that interact with uric acid (Figures 3A and 3B).

**Figure 3.**
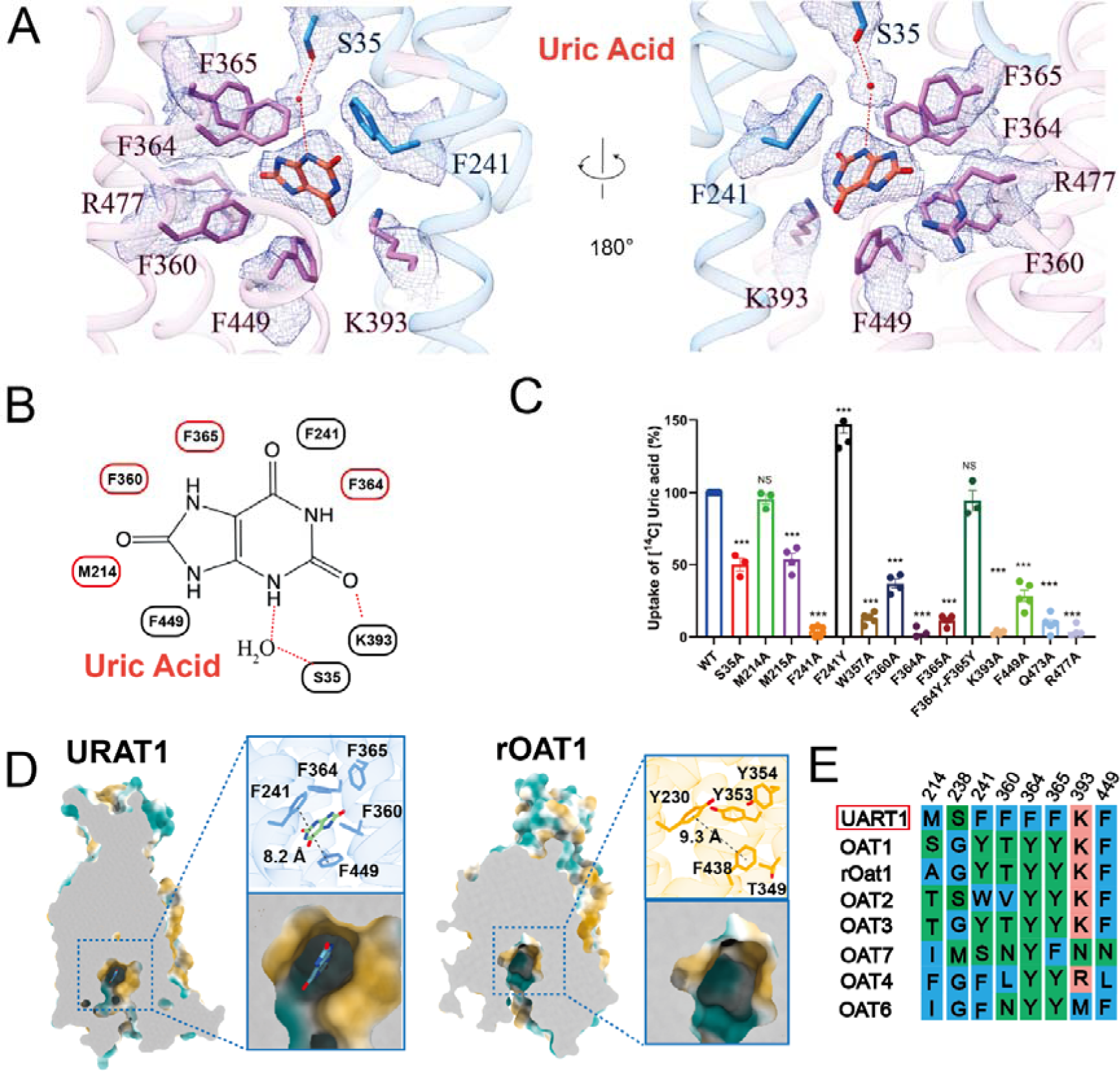
Recognition of uric acid by URAT1. **(A)**. The interaction network between uric acid and URAT1 residues is depicted. A red dot indicates a water molecule, hydrogen bonds are highlighted with red dashed lines, and the corresponding density maps of the residues and ligand are shown as a mesh. **(B)**. Schematic representation of the interactions between URAT1 and uric acid in 2D format. (**C**).^14^C-uric acid uptake activities of mutants of pocket-forming residues relative to the wild-type (WT) URAT1. Data are shown as “mean values ± S.E.M.”; Four independent replicates were performed. Data were analyzed by two-sided, one-way ANOVA Tukey’s test. *P < 0.05, **P < 0.01, ***P < 0.001. **(D)**. Cut-open surface view showing the substrate binding pocket in URAT1 and rOAT1. The distances from F241 to F449 in URAT1 and Y230 to F38 in rOAT1 are measured and marked. **(E)**. Sequence alignment of residues around the uric acid binding site in URAT1, OAT1, rOAT1, OAT2, OAT3, OAT7, OAT4, and OAT6.

The uric acid molecule primarily engages in π-stacking interactions with phenylalanine residues F241, F360, F364, F365, and F449. To validate the functional importance of these interactions, we performed alanine substitutions of these residues. These mutations significantly reduced the transport activity of URAT1 (Figure 3C), underscoring their crucial role in substrate recognition. Additionally, a polar interaction between uric acid and K393 further stabilizes the binding, as evidenced by the substantial decrease in transport activity upon K393A mutation. Interestingly, we observed that S35 in TM1 forms a water-mediated hydrogen bond with uric acid. Alanine substitution of S35 also resulted in a notable reduction in the transport activity of URAT1 (Figure 3C), highlighting the importance of this interaction.

The substrate-binding pocket of URAT1 exhibits unique features compared to other anion transporters and SLC22 family members. While transporters like OAT1 typically employ a combination of phenylalanine and tyrosine residues to create their substrate-binding aromatic cages, URAT1 exclusively utilizes phenylalanine residues (Figures 3D and 3E). This characteristic likely renders the binding pocket of URAT1 more hydrophobic than those of other transporters. Moreover, the binding pocket is relatively narrow, with an 8.2 Å separation between F241 and F449, compared to approximately 9.1 Å in rOAT1(Figure 3D). These features impose stringent selection criteria based on substrate polarity and size, contributing to the high substrate specificity observed for URAT1.

In contrast to the selectivity of URAT1, OAT1, OAT4, and OAT6 exhibit lower substrate specificity and can efficiently transport multiple substrates, including uric acid^22^. To explore the structural basis of this difference, we performed mutation studies. Intriguingly, mutating both F364 and F365 in URAT1 to tyrosine, as found in OAT1, OAT4, and OAT6, did not affect its transport activity for uric acid. However, the F241Y mutation did impact transport activity (Figure 3C), suggesting that while some structural variations are tolerated, certain key residues are critical for maintaining the specific uric acid transport capability of URAT1.

These findings provide crucial insights into the molecular basis of the selective uric acid transport of URAT1. The unique arrangement of aromatic residues in the binding pocket, coupled with its narrow dimensions, likely contributes to the high substrate specificity of URAT1 compared to other organic anion transporters. This structural information offers a foundation for the rational design of URAT1-targeted therapeutics and helps explain the crucial role of the transporter in uric acid homeostasis.

### Inhibition of URAT1 by benzbromarone and dotinurad

Building upon our structural insights into uric acid recognition by URAT1, we next investigated how this transporter is inhibited by uricosuric agents. We first focused on two drugs: benzbromarone, introduced in the 1970s^9^, and dotinurad, a newer agent derived from benzbromarone approved in Japan in 2020^17^. While both drugs enhance uric acid excretion, dotinurad offers more selective inhibition of uric acid reabsorption with fewer off-target effects.

We first evaluated the inhibitory effects of benzbromarone and dotinurad on URAT1-mediated ¹LJC-uric acid uptake, revealing ICLJLJ values of ∼200 nM and 8 nM, respectively (Figure 4A, and Table S2). The cryo-EM structures of URAT1 in complex with the two drugs, resolved at 3.2 Å and 3.6 Å respectively, revealed that both compounds occupy the central binding pocket of URAT1 at the uric acid binding site, stabilizing the transporter in an inward-open conformation. Despite their structural similarities, the two compounds exhibit distinct binding modes, which we categorized into Part A and Part B for clarity (Figure 4B).

**Figure 4.**
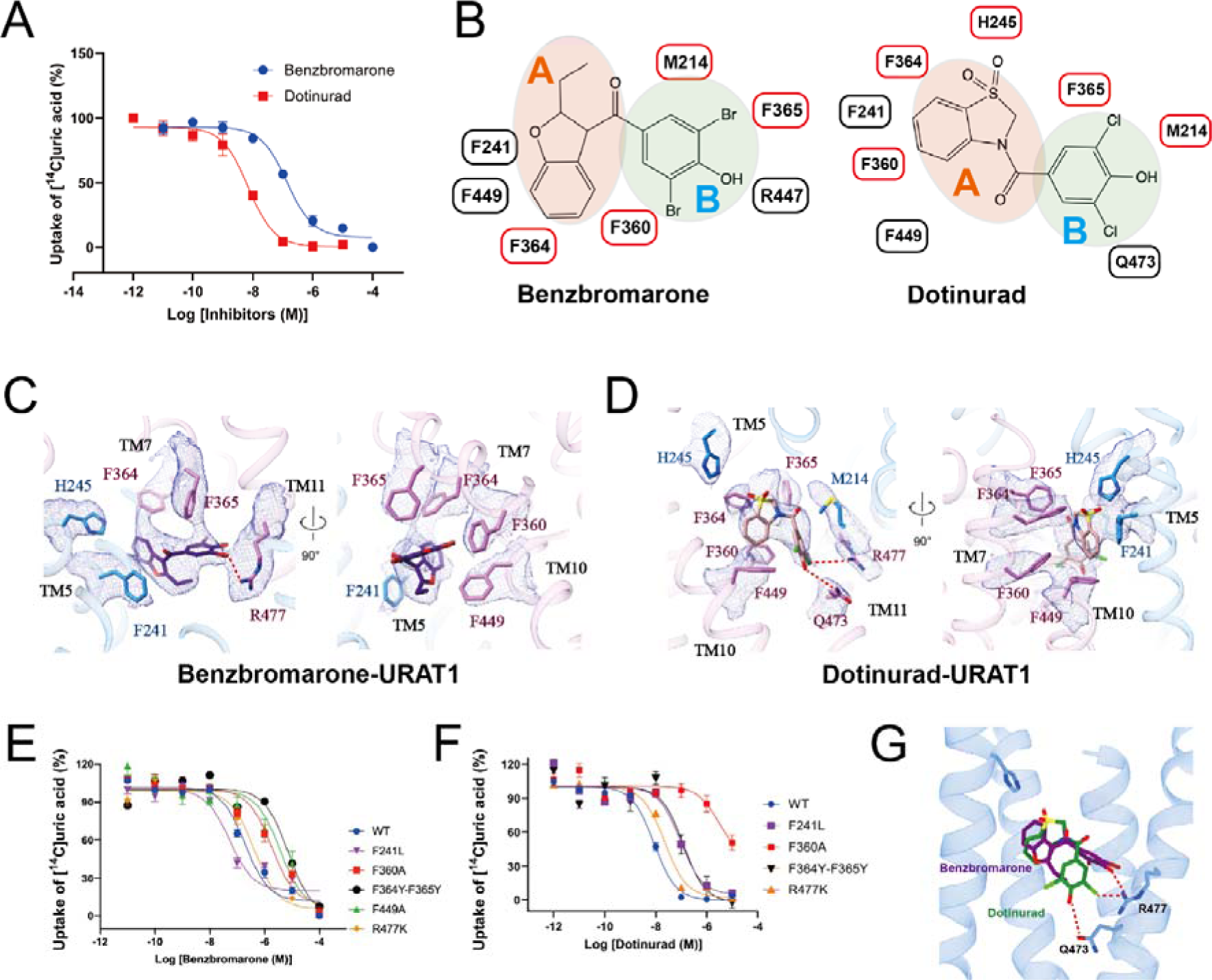
Inhibition of URAT1 by benzbromarone and dotinurad. **(A)**. Dose-dependent inhibition of ^14^C-uric acid uptake by Benzbromarone (blue) and Dotinurad (red) on WT URAT1. Data are shown as “mean values ± S.E.M.” Three independent replicates were performed, and all curves were fitted by nonlinear regression. **(B)**. Schematic representation of the interaction networks between Benzbromarone and Dotinurad with URAT1 residues, depicted in 2D format, including the division of ligand regions. **(C)**. The interaction network between benzbromarone and URAT1 residues is depicted. Hydrogen bonds are highlighted with red dashed lines, and the corresponding density maps of the residues and ligand are shown as a mesh. **(D)**. The interaction network between dotinurad and URAT1 residues is depicted. Hydrogen bonds are highlighted with red dashed lines, and the corresponding density maps of the residues and ligand are shown as a mesh. **(E)** and (**F**). Dose-dependent inhibition of ^14^C-uric acid uptake by Benzbromarone (E) and Dotinurad (F) in WT URAT1 and URAT1 mutants. (**G**). Structural alignment of Benzbromarone and Dotinurad in URAT1.

Part A of both molecules engages with the uric acid binding site, albeit in different poses. Benzbromarone primarily forms hydrophobic interactions with surrounding phenylalanine residues, including F241, F360, F364, F365, and F449 (Figures 4C and 4D). In contrast, dotinurad, while also interacting with F360, F365, and F449, forms an anion-π interaction with F241 and F364 via its sulfuryl group (Figures 4C and 4D). These interactions mirror the π-stacking interactions observed between uric acid and the phenylalanine residues in the substrate-bound structure. Alanine substitutions at F360 or F449 in URAT1 significantly diminish the inhibitory effectiveness of benzbromarone (Figure 4E, and Table S2). Additionally, substituting alanine at F360 markedly reduces the effectiveness of dotinurad inhibition (Figure 4F, and Table S2). These findings underscore the critical role of these residues in substrate recognition, as previously demonstrated.

Intriguingly, URAT1 demonstrates structural flexibility to accommodate the different geometries of benzbromarone and dotinurad. It creates a larger cavity between F241 and F360 for the bulkier benzbromarone (8.5 Å) compared to dotinurad (7.4 Å) (Figure S6). This adaptability is reflected in mutational studies: replacing F241 with the smaller leucine increased the potency of benzbromarone 4-folds but reduced the potency of dotinurad by 13-folds (Figures 4E and 4F; and Table S2). This difference in pocket size may explain higher potency of dotinurad in inhibiting URAT1 transport and highlights the importance of the dimensions of binding pocket in substrate specificity, as observed in the uric acid-bound structure.

Part B of the inhibitors interacts with a sub-pocket not engaged by uric acid. Benzbromarone extends into this area bordered by TM4, TM7, and TM11, interacting with M214, F360, and F365 (Figures 4C and 4D). Its bromine group engages in polar interactions with R477, and mutation of R477 to lysine slightly reduced the potency of benzbromarone (Figures 4E and 4F; and Table S2). In contrast, Part B of dotinurad rotates approximately 90 degrees relative to benzbromarone, maintaining interactions with M214, F360, and F365, while its hydroxyl group forms hydrogen bonds with both Q473 and R477 (Figure 4G). The different orientations of Part B in both inhibitors likely contributes to higher selectivity and potency of dotinurad compared to benzbromarone.

These distinct binding modes of benzbromarone and dotinurad within the URAT1 active site not only explain their different potencies but also provide valuable structural insights for the rational design and optimization of next-generation URAT1-targeted therapeutics for managing hyperuricemia and gout. The comparison between substrate and inhibitor binding mechanisms reveals how subtle structural differences can significantly impact binding affinity and specificity, underscoring the importance of detailed structural studies in drug development.

### Inhibition of URAT1 by lesinurad and verinurad

Building upon our analysis of benzbromarone and dotinurad, we expanded our investigation of URAT1 inhibition to include lesinurad, a uricosuric agent approved in 2015^23^, and its successor, verinurad. Verinurad was developed to address nephrotoxicity concerns of lesinurad, demonstrating higher activity and selectivity for URAT1 in Phase III clinical trials^24^.

Our assessment of URAT1-mediated ¹LJC-uric acid uptake inhibition revealed ICLJLJ values of ∼12 μM for lesinurad and 40 nM for verinurad (Figure 5A), indicating the higher potency of verinurad. Cryo-EM structures of URAT1 in complex with the two inhibitors, resolved at 3.4 Å and 3.2 Å respectively, unveiled distinct binding modes despite their similar chemical scaffolds. While our earlier analysis of benzbromarone and dotinurad revealed two distinct binding regions (Part A and Part B), our examination of lesinurad and verinurad led to a refined model with three key interaction regions within the URAT1 binding site: A, B, and C (Figure 5B). This expanded model provides a more comprehensive framework for understanding inhibitor binding while maintaining consistency with our earlier observations.

**Figure 5.**
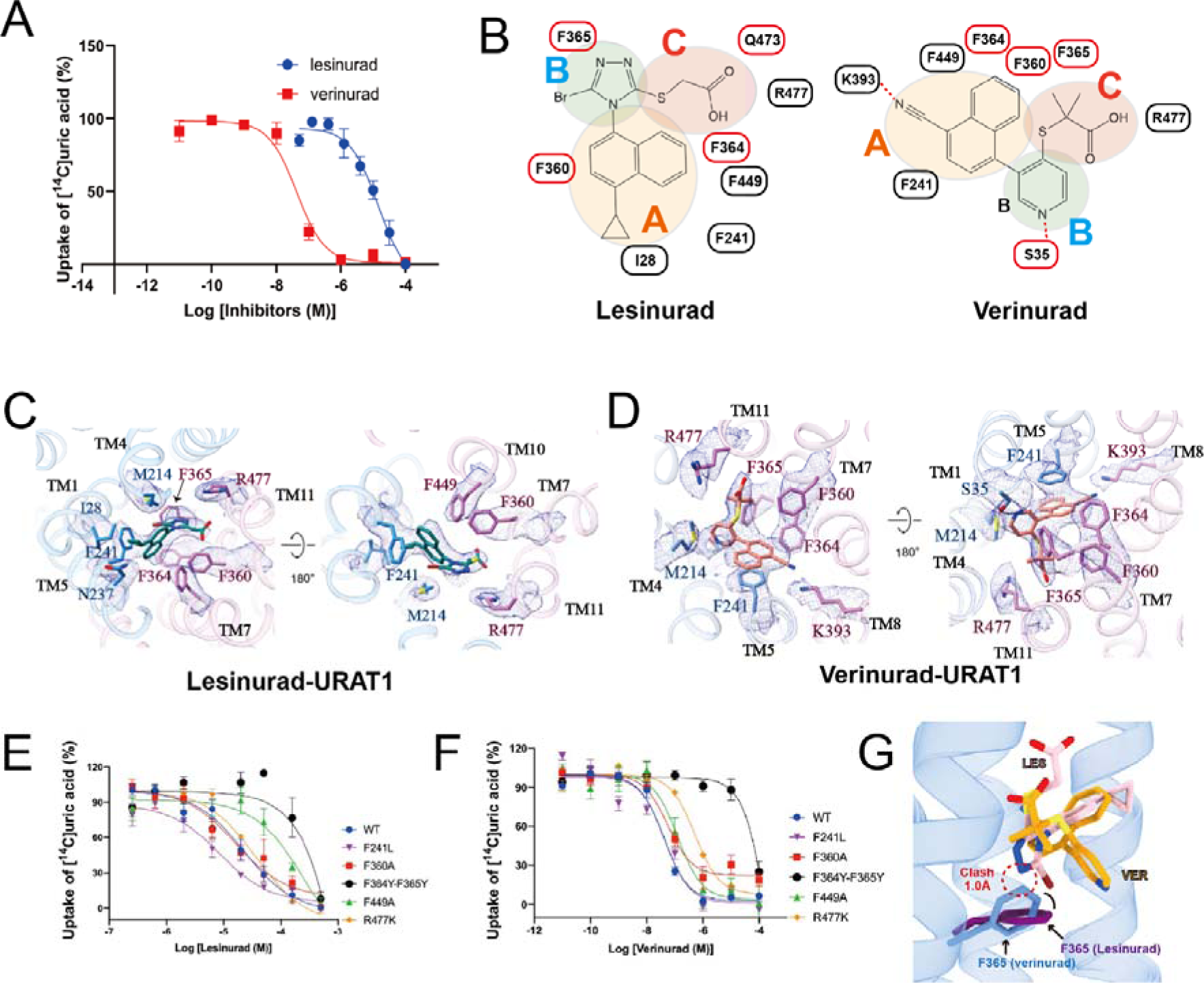
Inhibition of URAT1 by Lesinurad and Verinurad. **(A)**. Dose-dependent inhibition of ^14^C-uric acid uptake by Benzbromarone (blue) and Dotinurad (red) on WT URAT1. Data are shown as “mean values ± S.E.M.” Three independent replicates were performed, and all curves were fitted by nonlinear regression. **(B)**. Schematic representation of the interaction networks between lesinurad and verinurad with URAT1 residues, depicted in 2D format, including the division of ligand regions. **(C)**. The interaction network between lesinurad and URAT1 residues is depicted. Hydrogen bonds are highlighted with red dashed lines, and the corresponding density maps of the residues and ligand are shown as a mesh. **(D)**. The interaction network between verinurad and URAT1 residues is depicted. Hydrogen bonds are highlighted with red dashed lines, and the corresponding density maps of the residues and ligand are shown as a mesh. (**E**) and (**F**). Dose-dependent inhibition of ^14^C-uric acid uptake by lesinurad (E) and verinurad (**F**) in WT URAT1 and URAT1 mutants. (**G**). Structural alignment of lesinurad and verinurad in URAT1. The clash regions are circled.

Region A, analogous to Part A in our previous analysis, involves the naphthalene ring core of both inhibitors, which interacts with multiple phenylalanine residues in the uric acid binding pocket (Figures 5C and 5D). Alanine substitutions of F449 in URAT1 reduce the inhibitory effectiveness of both drugs (Figures 5E and 5F; and Table S2). The nitrile substituent in verinurad forms an electrostatic interaction with K393 in TM8, contributing to its potency (Figures 5B and 5D). In contrast, the hydrophobic cyclopropyl group of lesinurad interacts with I28 and F241 in a pocket formed by TM1 and TM4 (Figures 5B and 5C).

Region B, which corresponds to Part B in our earlier categorization, exhibits significant differences between the inhibitors, similar to the distinctions we observed between benzbromarone and dotinurad. The pyridine ring of verinurad forms hydrophobic stacking interactions with F364 and F365, while its nitrogen atom makes a polar contact with S35 (Figures 5B and 5D). In contrast, the bromo-substituted 1,2,4-triazole ring of lesinurad interacts with F364 and F365 on TM7, slightly displacing these residues compared to their positions in the verinurad-bound structure (Figure 5C). Interestingly, mutating the surrounding phenylalanine residues to smaller amino acids, such as F241L, increases the potency of lesinurad but has no effect on the potency of verinurad (Figures 5E, 5F; and Table S2). Additionally, mutations to larger amino acids markedly decrease inhibitory activity. Notably, changing F364 and F365 to tyrosine nearly abolishes the inhibitory effects of lesinurad (Figures 5E, 5F; and Table S2), underscoring the structural sensitivity of this region. Previous studies have demonstrated that introducing a carbon atom as a hinge between regions A and B in lesinurad enhances its potency by ∼50 folds, likely by avoiding clashes with F364 and F365^25^.

Region C represents an additional interaction area not prominent in our analysis of benzbromarone and dotinurad, highlighting the evolving complexity of newer URAT1 inhibitors. Both lesinurad and verinurad contain a terminal carboxyl group in this region. The lesinurad C region penetrates deeper, forming a polar interaction with Q473 (Figures 5B and 5C). Verinurad, positioned further from Q473, interacts only with R477. Notably, the two methyl substituents in region C of verinurad engage in hydrophobic interactions with F360 and F365, which could enhance its binding capability (Figures 5B and 5D).

This expanded structure-activity relationship model allows us to draw parallels between different classes of URAT1 inhibitors while accommodating the unique features of each compound. By refining our analysis from a two-part to a three-region model, we can more accurately describe the binding modes of these structurally diverse inhibitors, providing a unified framework for understanding URAT1 inhibition mechanisms. This comparative analysis provides crucial insights into the molecular interactions governing URAT1 inhibition by lesinurad and verinurad, explaining their differing potencies and highlighting key structural features for designing more effective and safer uricosuric agents.

### Mechanism of URAT1 transport

Having elucidated the binding modes of uric acid and various inhibitors, we next sought to understand the dynamic conformational changes underlying URAT1’s transport function. We resolved structures of URAT1 in three distinct conformational states: inward-open (3.3 Å), outward-open (4.1 Å), and occluded (4.7 Å) (Figures 6A, 6B, and 6C). Although the structures of URAT1 in outward-open and occluded states have modest resolutions, the TMD helical structures and the position of the bound uric acid can be resolved (Figures 6A, 6B, and 6C). Notably, the ECD of URAT1 is absent in the outward-open and occluded states, suggesting high flexibility in these conformations.

**Figure. 6.**
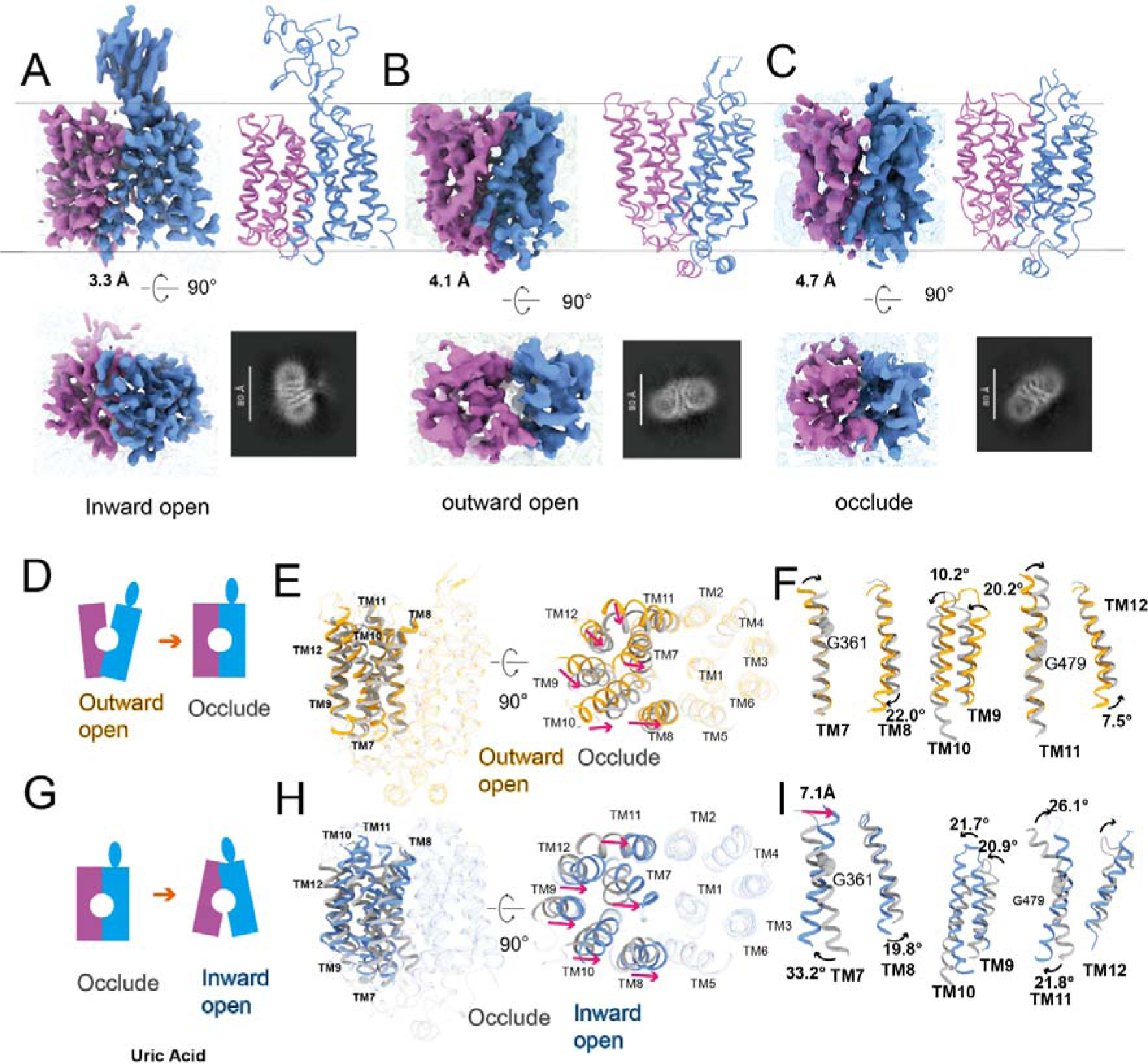
Mechanism of URAT1 transport. (**A**) to (**C**): 3D and 2D density maps and models of URAT1 in three distinct conformations: inward open state (**A**), outward open state (**B**), and occluded state (**C**). (**D**) A cartoon diagram illustrating the transition from the outward open to the occluded state of URAT1. (E) and (**F**) Structure alignment of the outward open state (yellow) and occluded state (gray) of URAT1, showing the overall structure alignment (**E**) and the alignment of transmembrane helices TM7/TM8, TM9/TM10, and TM11/TM12 (**F**). (**G**) A cartoon diagram depicting the transition from the occluded to the inward open state of URAT1. (**H**) and (**I**) Structure alignment from the occluded state (gray) to the inward open state (blue) of URAT1, including the overall structure alignment (**H**) and the alignment of transmembrane helices TM7/TM8, TM9/TM10, and TM11/TM12 (**I**). Red arrows indicate the direction of helix displacement, black arrows show the direction of helix rotation, and the rotation angles are also marked. The N-terminal domain (NTD) was used as a reference for structural alignments.

Analysis of these structures revealed that the NTD of URAT1 remains relatively stable across conformations, with an RMSD of 1.4 Å for Cα atoms between the outward-open and inward-open states. In contrast, the CTD undergoes significant rearrangements, exhibiting an RMSD of 4.5 Å for Cα atoms.

The transport cycle begins with the outward-open state, where a cavity formed by the extracellular termini of the NTD and CTD helices allows substrate entry. As URAT1 transitions to the occluded state, the extracellular termini of the CTD helices move inward, while the intracellular portions remain relatively static (Figures 6D and 6E). A key feature of this transition is the rotation of the extracellular terminus of TM7 toward TM1, moving 4.5 Å closer to the central cavity and significantly narrowing it (Figure. 6F). This movement hinges on Glycine 361 (G361) in TM7, explaining why the G361V mutation, found in patients with renal hypouricemia, disrupts transport (https://www.ncbi.nlm.nih.gov/clinvar/). Concurrently, TM8-TM12 in the CTD shift toward the NTD (Figures 6E and 6F).

The final transition to the inward-open state involves even more pronounced movements of the CTD helices (Figures 6G and 6H). the extracellular terminus of TM7 rotates an additional 20.3° toward TM1, forming polar interactions and nearly sealing the extracellular cavity. Simultaneously, its intracellular terminus moves 7.1 Å away from the NTD (Figure 6I). TM11 undergoes a substantial reconfiguration, with its extracellular and intracellular termini rotating 26.1° toward and 21.8° away from the NTD, respectively (Figure 6I). The importance of these movements is underscored by the G479R mutation in TM11, which is a loss-function mutation that is associated with renal hypouricemia (https://www.ncbi.nlm.nih.gov/clinvar/). TM8-TM10 and TM12 undergo similar shifts, collectively expanding the intracellular cavity for substrate release (Figure 6I).

These structural insights provide a comprehensive framework for understanding the transport mechanism of URAT1, revealing how coordinated conformational changes facilitate substrate translocation (Figure 7A). Importantly, the inward-open state can be stabilized by inhibitors, offering a structural explanation for their mode of action and providing valuable information for future drug design efforts (Figure 7B).

**Figure 7.**
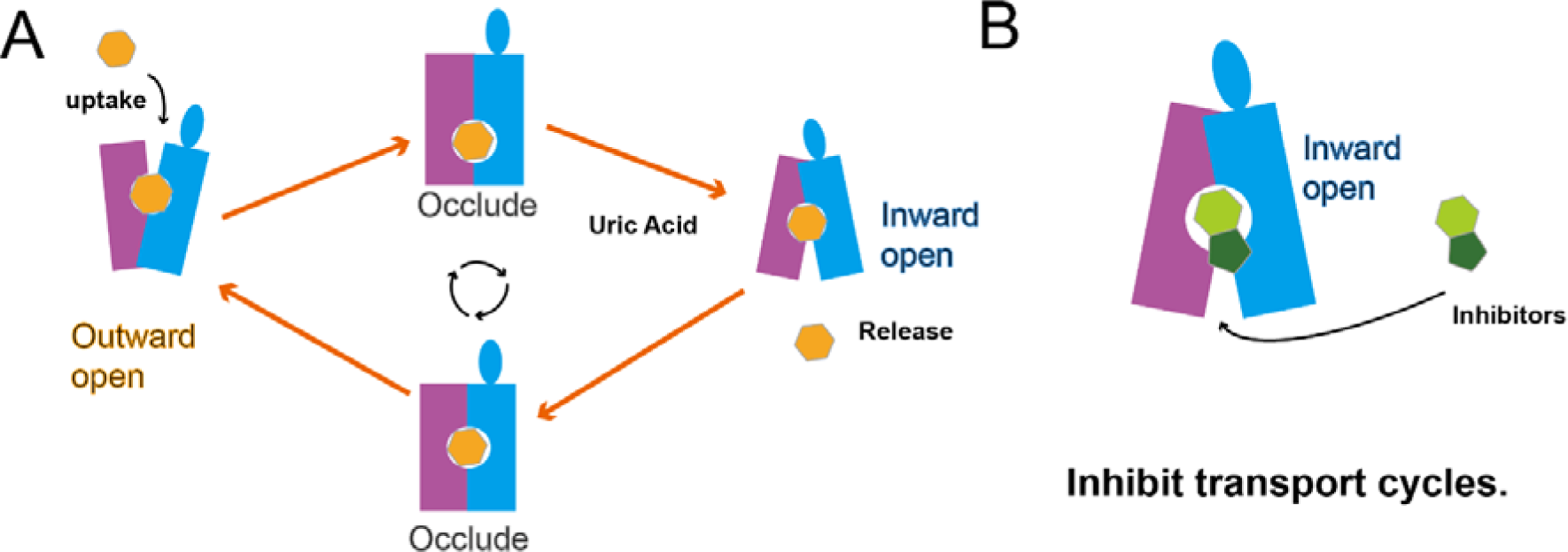
URAT1 transport cycle and mechanism of inhibition by anti-Gout drugs. (**A**) Schematic diagram illustrating the transport of uric acid by URAT1. (**B**) Schematic diagram depicting the inhibition of URAT1 by anti-gout drugs.

## Discussion

In this paper, we have overcome technical challenges in expressing and purifying the human URAT1 for cryo-EM studies. The structural elucidation of URAT1 in complex with its substrate and various inhibitors provides important insights into the molecular mechanisms of urate transport and inhibition. Our study reveals that the substrate binding pocket of URAT1 is characterized by an aromatic cage composed exclusively of phenylalanine residues, a feature that distinguishes it from other organic anion transporters. This unique arrangement explains the high specificity of URAT1 for uric acid and provides a structural basis for the development of selective inhibitors. The observed π-stacking interactions between uric acid and residues F241, F360, F364, F365, and F449, along with the polar interaction with K393, highlight the critical residues involved in substrate recognition. These findings not only elucidate the molecular basis of the substrate specificity of URAT1 but also offer valuable targets for structure-guided drug design.

The structures of URAT1 bound to four clinically relevant inhibitors - benzbromarone, dotinurad, lesinurad, and verinurad - reveal distinct binding modes that explain their varying potencies and selectivity. Despite targeting the same pocket, these inhibitors exhibit different interaction patterns with URAT1. The comparison between benzbromarone and dotinurad is particularly informative, demonstrating how subtle differences in binding pocket size can significantly impact inhibitor potency. The higher potency of dotinurad, attributed to its more compact binding mode, provides a clear strategy for enhancing drug potency. Furthermore, the unique interactions observed with lesinurad and verinurad, such as the nitrile group of verinurad that forms an electrostatic interaction with K393, could offer additional avenues for optimizing inhibitor design.

Our resolution of URAT1 structures in three distinct conformational states - outward-open, occluded, and inward-open - provides a comprehensive view of the transport mechanism. The observed conformational changes, particularly in the C-terminal domain, reveal how URAT1 facilitates substrate translocation. The identification of key residues such as G361 and G479, which act as hinges for these conformational shifts, explains the impact of disease-associated mutations and offers potential targets for allosteric modulation of URAT1 function. The G361V and G479R mutations, identified in patients with renal hypouricemia, are now understood to disrupt critical conformational changes necessary for the transport function of URAT1. This knowledge not only explains the pathophysiology of these mutations but also suggests potential approaches for personalized treatment strategies in gout and related disorders. These insights enhance our understanding of the physiological role of URAT1 and suggest new strategies for therapeutic intervention.

The structural flexibility of the binding pocket of URAT1, as evidenced by its ability to accommodate diverse inhibitors, has significant implications for drug design. The observed adaptability of residues like F241 in response to different inhibitors suggests that rational modifications of inhibitor structure could exploit this flexibility to enhance binding affinity and selectivity. Moreover, the identification of unique residues in URAT1, such as Q473, which contributes to the higher selectivity of newer inhibitors like dotinurad, provides a clear direction for developing more specific URAT1-targeted therapeutics.

The potential impact of URAT1 inhibitors in gout treatment can be paralleled with the remarkable success of SGLT2 inhibitors in diabetes management. SGLT2 inhibitors have revolutionized diabetes treatment by targeting glucose reabsorption in the kidneys, effectively lowering blood sugar levels while offering additional cardiovascular and renal benefits. For instance, drugs like empagliflozin (Jardiance), dapagliflozin (Farxiga), and canagliflozin (Invokana) have shown significant reductions in HbA1c levels, body weight, and blood pressure in type 2 diabetes patients^26–28^. Moreover, these drugs have demonstrated cardiovascular benefits, with empagliflozin showing a 38% relative risk reduction in cardiovascular death in high-risk patients^26,27^. Similarly, URAT1 inhibitors have the potential to transform gout treatment by enhancing uric acid excretion. The structural insights provided by our study offer a comparable opportunity for the rational design and optimization of URAT1 inhibitors, potentially leading to a new generation of therapeutics with improved efficacy and reduced side effects.

In conclusion, this comprehensive structural study of URAT1 marks a significant advance in our understanding of urate transport and its inhibition. By providing a detailed structural framework for the function and inhibition of URAT1, our work paves the way for the rational design of next-generation therapeutics for hyperuricemia and gout. The insights gained from this study could have the potential to guide the development of more effective and selective URAT1 inhibitors, ultimately leading to improved treatment options for millions of patients worldwide affected by gout and related conditions.

## Materials and Methods

### Expression and purification of URAT1

The human wild-type URAT1 gene was cloned into a pEG-BacMam vector with an N-terminal Flag tag. The construct was confirmed by DNA sequencing. The Bac-to-Bac baculovirus expression system was used to produce recombinant baculovirus in Spodoptera frugiperda Sf9 insect cells (Invitrogen), and the P2 viruses were used to infect HEK293E cells for protein expression. 10 liters of HEK293E cells were cultured in SMM 293-TII Expression Medium (Sino Biologica) at 37°C with 5% CO2. Following a 10-hour transfection period, 10 mM sodium butyrate was added to enhance protein expression. The cells were cultured for an additional 48 hours at 37°C, then harvested by centrifugation at 2,000g for 15 minutes and stored at −80°C for purification.

The cells were thawed at 37°C, resuspended in a lysis buffer containing 20 mM HEPES (pH 7.5), 150 mM NaCl, 10% glycerol, and an EDTA-free protease inhibitor cocktail (TargetMol), and lysed mechanically. 0.5% lauryl maltose neopentyl glycol (LMNG, Anatrace) and 0.1% cholesterol hemisuccinate (CHS, Anatrace) were added to the lysate, and solubilization was carried out for 2 hours at 4°C. After centrifugation, the supernatant was incubated with anti-DYKDDDDK G1 Affinity Resin (GenScript) for 3 hours at 4°C. The resin was then packed and washed with 30 column volumes of 20 mM HEPES pH 7.4, 150 mM NaCl, 0.01% (w/v) LMNG, 0.002% CHS, and 10 μM ligand. The complex sample was eluted in a buffer containing 20 mM HEPES pH 7.4, 150 mM NaCl, 0.01% (w/v) LMNG, 0.002% CHS, 10 μM ligand, and 0.2 mg/ml FLAG peptide (GenScript). The complex fractions were concentrated using a 100-kDa molecular weight cut-off (MWCO) Millipore concentrator for further purification.

The complex was then subjected to size-exclusion chromatography on a Superdex 6 Increase 10/300 GL column (GE Healthcare) pre-equilibrated with a size buffer containing 20 mM HEPES pH 7.4, 150 mM NaCl, 0.00075% (w/v) LMNG, 0.00025% (w/v) GDN (Anatrace), and 0.00015% CHS.

To prepare URAT1 samples bound to verinurad, lesinurad, benzbromarone, dotinurad, or uric acid, 10 μM of the corresponding inhibitor or 1 mM for substrate was added throughout the purification buffer system. Before preparing Cryo-EM grids, an additional 100 μM of the respective inhibitor was supplemented.

### Cryo-EM Data Collection

Cryo-EM grids were prepared using a Vitrobot Mark IV (FEI) at 6°C and 100% humidity. A 3 μL aliquot of the sample was applied to glow-discharged gold R1.2/1.3 holey carbon grids. The sample was allowed to incubate on the grids for 10 seconds before being blotted for 4.5 seconds (double-sided, blot force 1) and then rapidly frozen in liquid ethane.

All datasets were acquired using a Titan Krios microscope fitted with a Falcon 4i direct electron detection camera operating at 300 kV and a magnification of 165,000x, resulting in a pixel size of 0.73 Å. Image acquisition was managed with EPU Software (FEI Eindhoven, Netherlands). A total of 36 frames were captured per movie, accumulating to a total electron dose of 50 e-/Å² over an exposure time of 2.5 seconds.

### Cryo-EM image processing

MotionCor2 was used to perform frame-based motion correction and generate drift-corrected micrographs for further processing^29,30^. All subsequent steps, including contrast transfer function (CTF) estimation, particle picking and extraction, two-dimensional (2D) classification, ab initio reconstruction, heterogeneous refinement, non-uniform refinement, local refinement, and local resolution estimation, were performed using cryoSPARC^31^.

For the URAT1-verinurad dataset, 7,883 dose-weighted micrographs were imported into cryoSPARC, and CTF parameters were estimated using patch-CTF (Figure S2, and Table S1). Initially, 2,338,086 particles were picked using the blob picker from the full set of micrographs and extracted with a pixel size of 1.46 Å. After multiple rounds of 2D classification, 49,249 particles were selected and extracted with a pixel size of 0.73 Å for further ab initio reconstruction, yielding a 4.7 Å map through non-uniform refinement.

Subsequently, 20 2D templates were generated from the 3D map, and 11,742,200 particles were picked using the template picker from an enlarged set of micrographs and extracted with a pixel size of 1.46 Å. After one round of 2D classification, the selected particles underwent several rounds of multi-reference guided 3D classification. The good particles were extracted with a pixel size of 0.73 fGOfor further ab initio reconstruction, followed by non-uniform refinement and local refinement, generating a 3.2 Å map. A similar processing workflow was followed for the URAT1-lesinurad dataset, yielding a 3.5 Å map (Figure S2), the URAT1-benzbromarone dataset, yielding a 3.3 Å map, and the URAT1-dotinurad dataset, yielding a 3.6 Å map (Figure S3; and Table S1).

For the URAT1-uric acid dataset, 22,732 dose-weighted micrographs were imported into cryoSPARC, and the CTF parameters were estimated using patch-CTF (Figure S4, and Table S1). 15,643,611 particles were picked using the blob picker and template picker from the full set of micrographs and extracted with a pixel size of 1.46 Å. After multiple rounds of 2D classification, 1,036,225 particles were selected and extracted with a pixel size of 0.73 Å for further 2D classification. The good 2D classes were separated into two parts based on the presence or absence of ECD features. Particles from 2D classes with ECD features were used to generate an inward-open structure by ab initio reconstruction, whereas particles from 2D classes without ECD features were used to generate an outward-open and an occluded structure by ab initio reconstruction.

Multi-reference guided 3D classification was performed using these initial maps and the particles selected from the first round of 2D classification. Subsequently, ab initio reconstruction, non-uniform refinement, and local refinement were carried out, yielding the inward-open, outward-open, and occluded structures with resolutions of 3.1 Å, 4.1 Å, and 4.7 Å, respectively (Figure S4, and Table S1).

### Model building

A predicted URAT1 structure from Alphafold2 was used as the starting reference model for receptor building^32^. All models were fitted into the EM density map using UCSF Chimera^33^ followed by iterative rounds of manual adjustment and automated rebuilding in COOT^34^ and PHENIX^35^, respectively. The model was finalized by rebuilding in ISOLDE^36^ followed by refinement in PHENIX with torsion-angle restraints to the input model. The final model statistics were validated using Comprehensive validation (cryo-EM) in PHENIX^35^ and provided in the Supplementary Table 1. All structural figures were prepared using Chimera^33^, Chimera X^37^, and PyMOL (Schrödinger, LLC.).

### 14C-Uric acid uptake assay

The radiolabelled substrate uptake assays were performed using ^14^C-uric acid (American Radiolabeled Chemicals) to determine the transporter activity of different drugs towards wild-type and mutant hURAT1. HEK293 cells were cultured in Dulbecco’s modified Eagle’s medium (DMEM) supplemented with 10% FBS and 1% Sodium pyruvate and seeded at a density of 30,000 cells per well in Isoplate-96 plates (Perkin Elmer). Twenty-four hours after transfection with the wild-type or mutant hURAT1, the HEK293 cells were washed twice and incubated with assay buffer consisting of 25 mM HEPES, 125 mM sodium gluconate, 4.8 mM potassium gluconate, 1.2 mM monobasic potassium phosphate, 1.2 mM magnesium sulfate, 1.3 mM calcium gluconate and 5.6 mM glucose. For the single-point uptake assay, ^14^C-uric acid (50 μM) was added and reacted with the cells in assay buffer at room temperature for 15 min. The cells were then washed three times with wash buffer consisting of 25 mM HEPES and 125 mM sodium gluconate and lysed with 50 μL of 0.1 M NaOH. The radioactivity was subsequently counted (counts per minute, CPM) in a scintillation counter (MicroBeta 2 Plate Counter, Perkin Elmer) using a scintillation cocktail (OptiPhase SuperMix, Perkin Elmer). For the uptake assay with the hURAT1 inhibitors, compounds were serially diluted into assay buffer and added to the cells for 15 minutes and then ^14^C-uric acid (50 μM) were added and incubated for a further 15 min. Cells were then washed and lysed for liquid scintillation counting. Data were analysed by nonlinear regression using GraphPad Prism v.9.

The radiolabeled substrate uptake assays were performed using ^14^C-uric acid (American Radiolabeled Chemicals) to determine the transporter activity of different drugs towards wild-type and mutant human URAT1. HEK293 cells were cultured in Dulbecco’s modified Eagle’s medium (DMEM) supplemented with 10% fetal bovine serum (FBS) and 1% sodium pyruvate. The cells were seeded at a density of 30,000 cells per well in Isoplate-96 plates (Perkin Elmer). Twenty-four hours after transfection with the wild-type or mutant URAT1 plasmid, the cells were washed twice with an assay buffer consisting of 25 mM HEPES, 125 mM sodium gluconate, 4.8 mM potassium gluconate, 1.2 mM potassium phosphate monobasic, 1.2 mM magnesium sulfate, 1.3 mM calcium gluconate, and 5.6 mM glucose.

For the single-point uptake assay, ^14^C-uric acid (50 μM) was added to the cells in the assay buffer and incubated at room temperature for 15 minutes. Subsequently, the cells were washed three times with a wash buffer containing 25 mM HEPES and 125 mM sodium gluconate, and then lysed with 50 μL of 0.1 M NaOH. The radioactivity was measured as counts per minute (CPM) using a scintillation counter (MicroBeta 2 Plate Counter, Perkin Elmer) with a scintillation cocktail (OptiPhase SuperMix, Perkin Elmer).

For the uptake assay with hURAT1 inhibitors, the compounds were serially diluted in the assay buffer and added to the cells for 15 minutes. Then, ^14^C-uric acid (50 μM) was added, and the mixture was incubated for an additional 15 minutes. After incubation, the cells were washed and lysed for liquid scintillation counting. The data were analyzed by nonlinear regression using GraphPad Prism v.9.

### Statistics

Data from all functional studies were processed using GraphPad Prism version 9.0 (Graphpad Software Inc.), with results presented as mean values ± SEM, based on a minimum of three independent experiments, each conducted in triplicate. Statistical significance was assessed using a two-sided, unpaired t-test, where a p-value of less than 0.05 was deemed to indicate significant differences.

## Supporting information

Supplementary_imformation

## Acknowledgements

The cryo-EM data were collected at the Shanghai Advanced Center for Electron Microscopy, Shanghai Institute of Materia Medica, Chinese Academy of Sciences. We thank K.W., S.Z., and S.L. for helping with cryo-EM data collection. This work was partially supported by the National Key R&D Program of China (2022YFC2703105 to H.E.X.); CAS Strategic Priority Research Program (XDB37030103 to H.E.X.); Shanghai Municipal Science and Technology Major Project (2019SHZDZX02 to H.E.X.); Shanghai Municipal Science and Technology Major Project (H.E.X.); The National Natural Science Foundation of China (32130022 to H.E.X., 32301016 to C.W.,32171187 to Y.J., 82121005 to Y.J. and H.E.X. 82273985 to D.H.Y. 82121005 to D.H.Y.); the National Key Basic Research Program of China 2023YFA1800804 (D.H.Y.); Shanghai Municipality Science and Technology Development Fund 21JC1401600 (D.H.Y.) and Program of Shanghai Academic/Technology Research Leader 23XD1400900 (D.H.Y.); Hainan Provincial Major Science and Technology Project ZDKJ2021028 (D.H.Y.).

## Author contributions

C.W. and S.J designed the expression constructs, purified the protein complex supervised by H.E.X. and Y.J.. C.W. prepared the grids and performed the cryo-EM data processing and model building. H.Z., X.Y., and Y.X. contributed to data processing and model building. J.J.W. and C.W. constructed all the mutated plasmids, C.Z. and A.D. performed functional studies supervised by D.Y. and M.W.. H.E.X., and C.W. analyzed the structures. J.J.W. and C.W. prepared the figures and initial manuscript. Q.Y., and W.H. for performing cryo-EM data collection and Motion correlation. J.J.W., J.X. and X.H. helped with experiments. H.E.X. and C.W. conceived the project and initiated collaborations with D.Y. and Y.J., All authors discussed and commented on the manuscript. H.E.X. revised the paper, H.E.X. supervised the project. H.E.X., and C.W. wrote the manuscript with input from all authors.

## Competing interests

C.W., C.Z., S.J., J.J.W., A.D., J.X., H.Z., X.H., X.Y., Q.Y., W.H., Y.X., M.W., Y.J., D.Y., and H.E.X. have declared no competing interest.

## Notes

### Competing Interest Statement

The authors have declared no competing interest.

