## Supplementary_imformation for "Molecular mechanisms of uric acid transport by the native human URAT1 and its inhibition by anti-gout drugs"

Canrong Wu *et al.*

**This PDF file includes:**

Figs. S1 to S6
Table S1 to S2

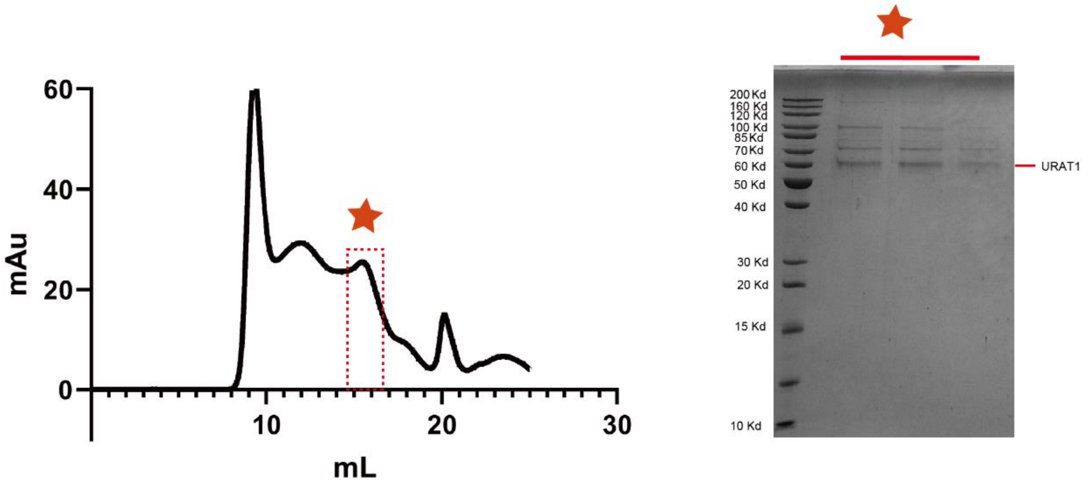


**Supplementary Figure 1. Representative size-exclusion chromatography and SDS-PAGE analysis of native hURAT1.**

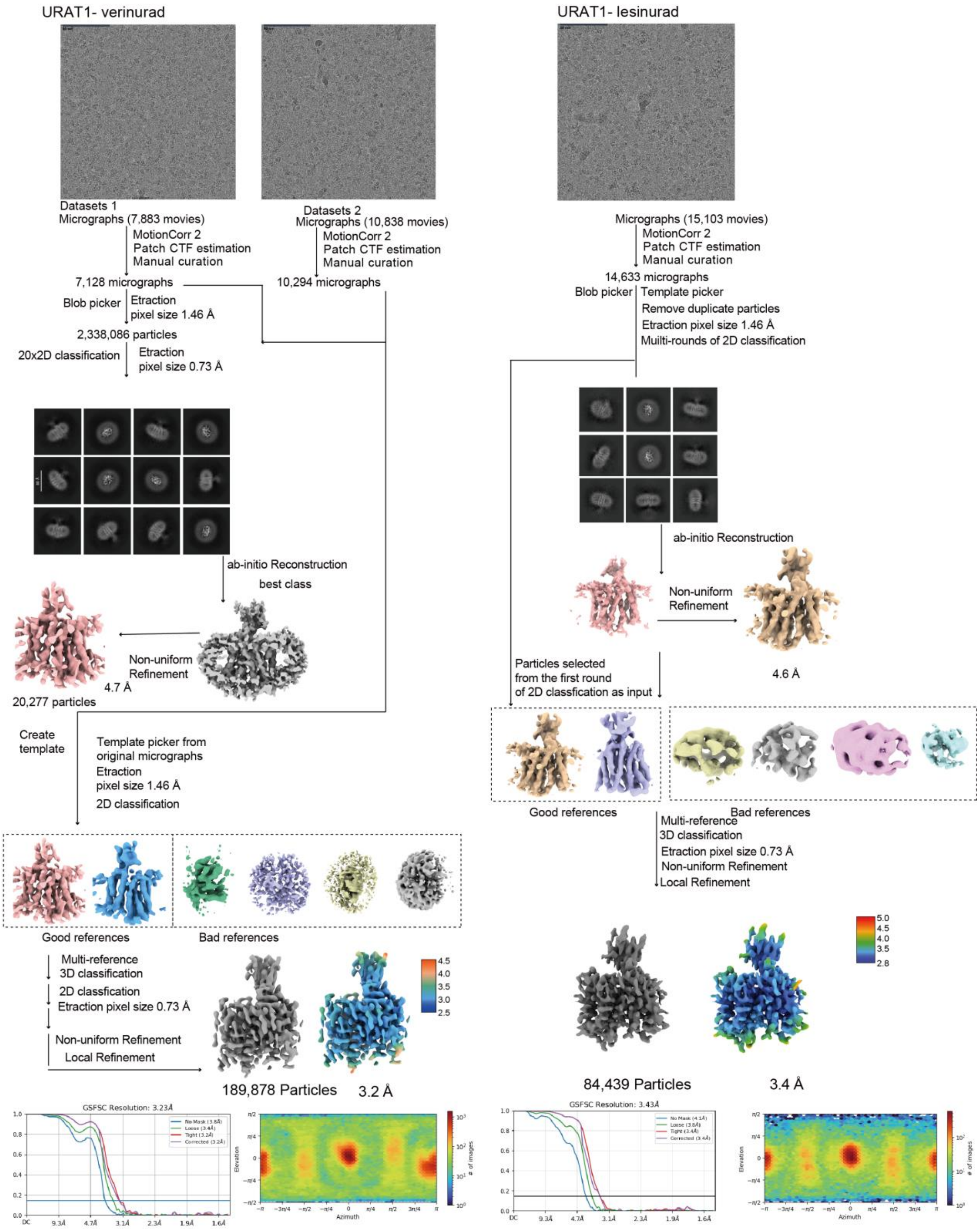


**Supplementary Figure 2.** A flow-chart of the cryo-EM data process of the URAT1- verinurad and URAT1-lesinurad.

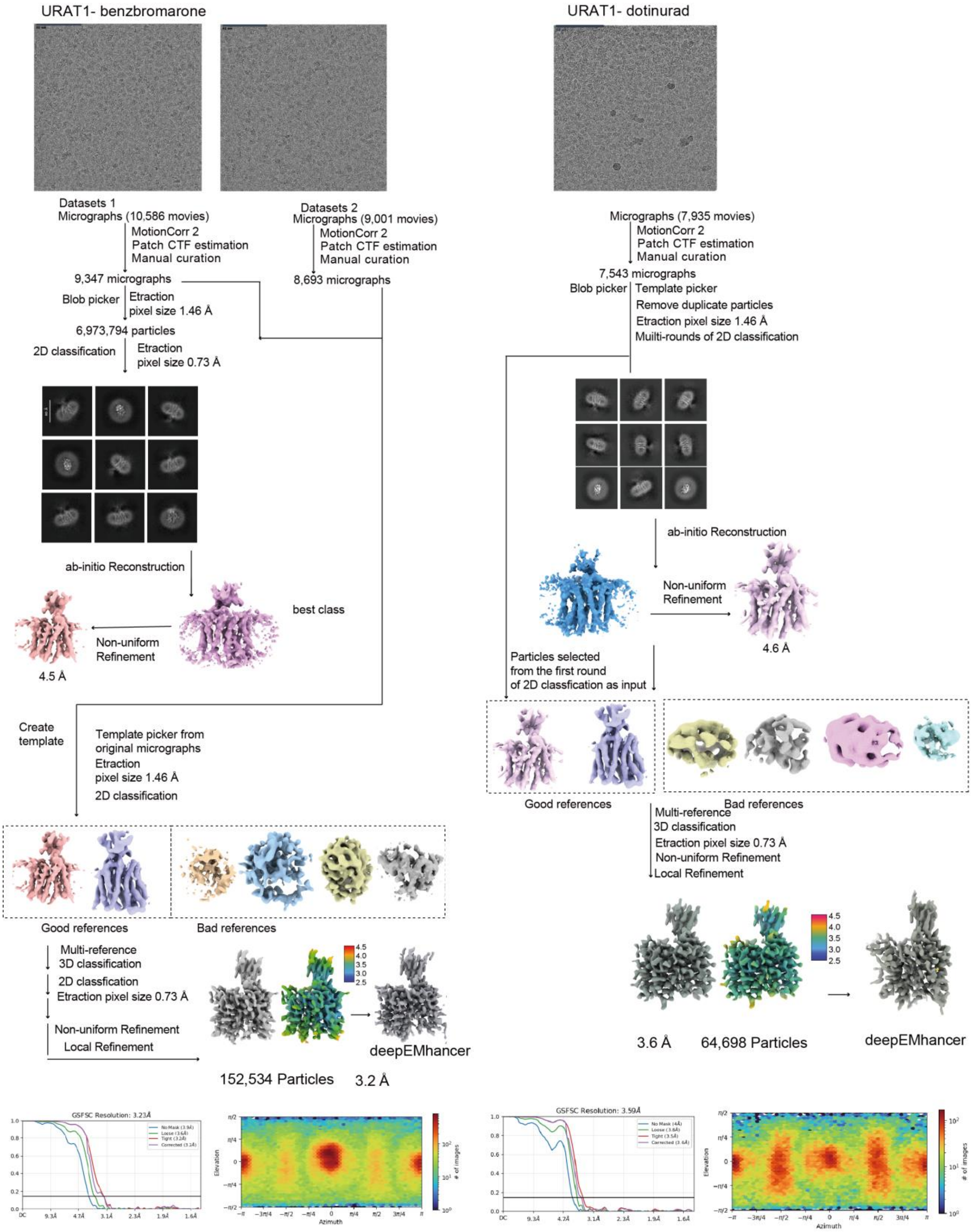


**Supplementary Figure 3.** A flow-chart of the cryo-EM data process of the URAT1-benzbromarone and URAT1-dotinurad.

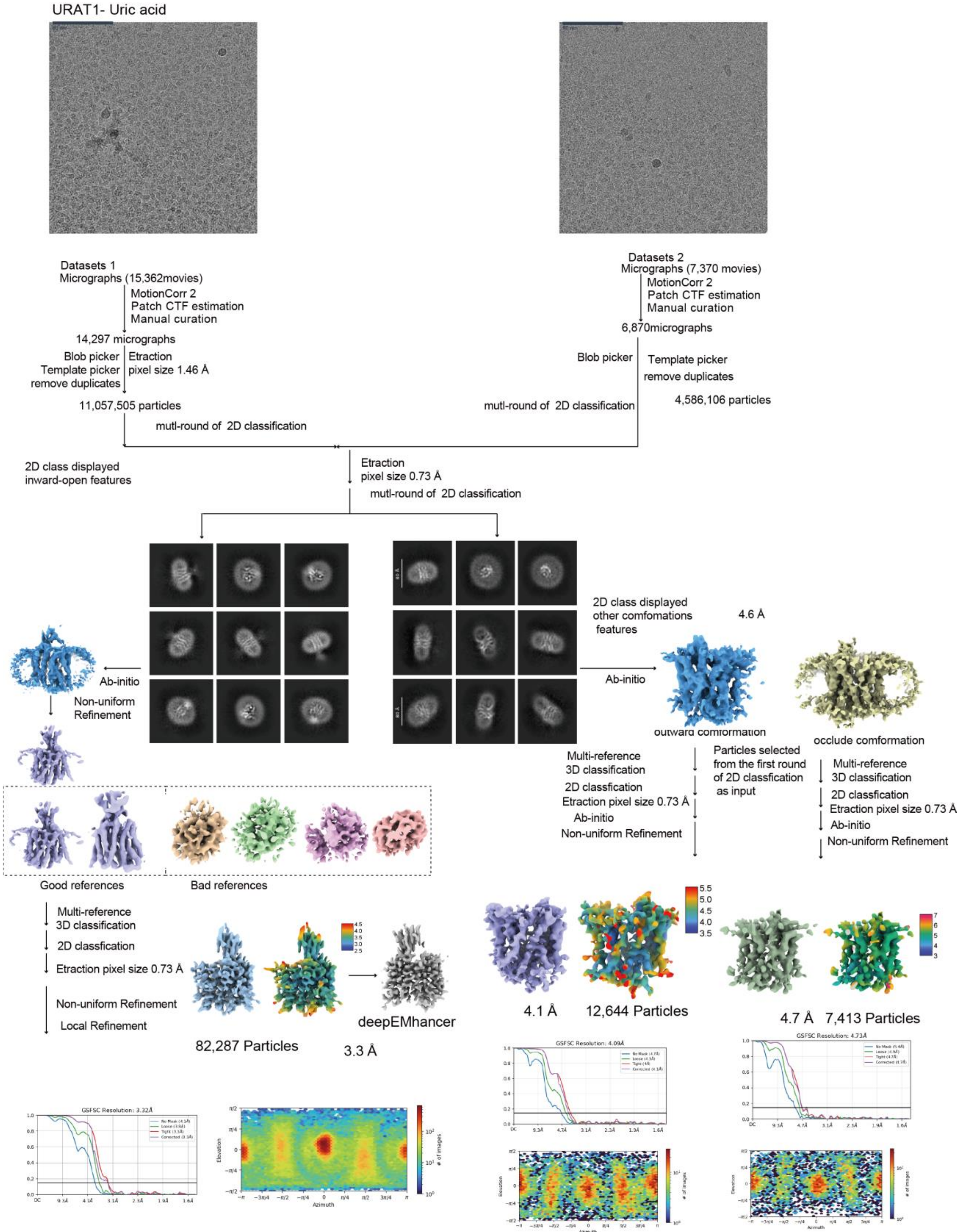


**Supplementary Figure 4.** A flow-chart of the cryo-EM data process of the URAT1-uric acid.

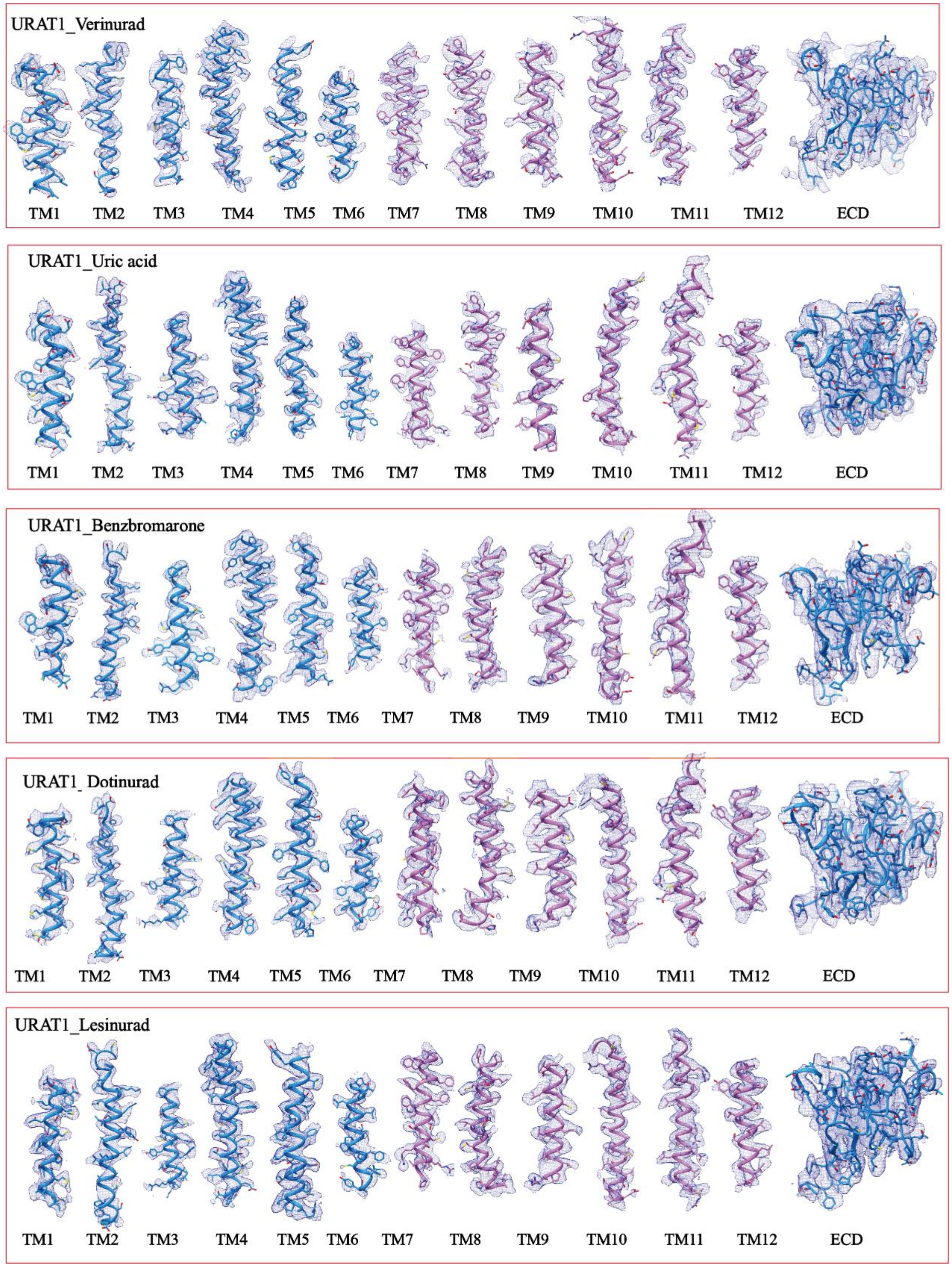


**Supplementary Figure 5.** The cryo-EM density map of the representative regions of the five URAT1 structures.

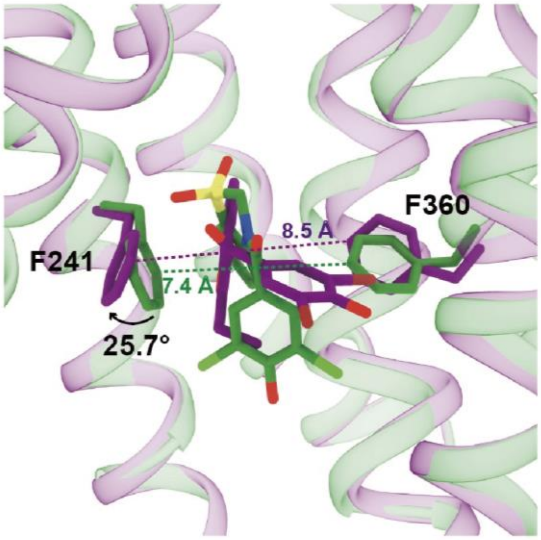


**Supplementary Figure 6. Different binding poses of benzbromarone and dotinurad in the URAT1 binding pocket.**

**Supplementary information, Table S1**

**Table S1.** Cryo-EM data collection, model refinement and validation statistics.

| | URAT1 verinurad | URAT1 dotinurad | URAT1 benzbromarone | URAT1 lesinurad | URAT1 Uric acid |
|---|---|---|---|---|---|
| **Data collection and processing** | | | | | |
| Detector | Falcon4 | Falcon4 | Falcon4 | Falcon4 | Falcon4 |
| Magnification | 165,000 | 165,000 | 165,000 | 165,000 | 165,000 |
| Voltage (kV) | 300 | 300 | 300 | 300 | 300 |
| Electron exposure ($e^{-}/Å^{2}$) | 50 | 50 | 50 | 50 | 50 |
| Defocus range (μm) | -1.0~-3.0 | -1.0~-3.0 | -1.0~-3.0 | -1.0~-3.0 | -1.0~-3.0 |
| Pixel size (Å) | 0.73 | 0.73 | 0.73 | 0.73 | 0.73 |
| Symmetry imposed | C1 | C1 | C1 | C1 | C1 |
| Initial particle projections (no.) | 11,742,200 | 5,792,635 | 10,776,212 | 8,668,180 | 15,643,611 |
| Final particle projections (no.) | 189,878 | 64,698 | 152,534 | 84,439 | 82,287 |
| Map resolution (Å) | 3.2 | 3.6 | 3.2 | 3.5 | 3.3 |
| Map resolution range (Å) | 2.5-4.5 | 2.5-4.5 | 2.5-4.5 | 2.8-4.5 | 2.5-4.5 |
| FSC threshold | 0.143 | 0.143 | 0.143 | 0.143 | 0.143 |
| **Model Refinement** | | | | | |
| Refinement package | PHENIX-1.17.1-3660 | PHENIX-1.17.1-3660 | PHENIX-1.17.1-3660 | PHENIX-1.17.1-3660 | PHENIX-1.17.1-3660 |
| Real or reciprocal space | Real space | Real space | Real space | Real space | |
| Model-Map CC (mask) | 0.66 | 0.69 | 0.68 | 0.65 | 0.66 |
| Model resolution (Å) | 3.3 | 3.7 | 3.4 | 3.5 | 3.5 |
| FSC threshold | 0.143 | 0.143 | 0.143 | 0.143 | 0.143 |
| B factors ($Å^{2}$, min/max/mean value) | | | | | |
| Protein residues | 39.63/134.22/67.53 | 38.52/146.58/70.54 | 63.50/142.74/93.31 | 65.83/161.05/94.37 | 67.61/136.11/93.07 |
| Ligands | 56.88/56.88/56.88 | 63.23/63.23/63.23 | 102.29/102.29/102.29 | 97.38/97.38/97.38 | 90.05/90.05/90.05 |
| **Model composition** | | | | | |
| Non-hydrogen atoms | 3,324 | 3,321 | 3,322 | 3,463 | 3,324 |
| Protein residues | 435 | 435 | 435 | 440 | 435 |
| R.m.s. deviations | | | | | |
| Bond lengths (Å) | 0.001 | 0.002 | 0.002 | 0.002 | 0.001 |
| Bond angles (°) | 0.365 | 0.553 | 0.506 | 0.506 | 0.365 |
| **Validation** | | | | | |
| MolProbity score | 1.97 | 1.45 | 1.60 | 1.52 | 1.97 |
| Clashscore | 5.55 | 6.88 | 4.93 | 5.68 | 30.55 |
| Rotamer outliers (%) | 0.00 | 0.34 | 0.00 | 0.00 | |
| Ramachandran plot | | | | | |
| Favored (%) | 98.12 | 97.65 | 97.88 | 97.88 | 98.59 |
| Allowed (%) | 1.88 | 2.35 | 2.12 | 1.74 | 0.87 |
| Disallowed (%) | 0.00 | 0.00 | 0.00 | 0.00 | 0.00 |
| **Data availability** | | | | | |
| EMDB entry | EMD-61401 | EMD-61404 | EMD-61403 | EMD-61402 | EMD-61399 |
| PDB entry | 9JDY | 9JE1 | 9JE0 | 9JDZ | 9JDV |

**Supplementary Table 2 Effects of hURAT1 mutations on the transporter activity of different drugs.** The radiolabeled substrate-uptake assay of four compounds to the wild-type (WT) and mutated hURAT1 was performed in HEK293 cells with $^{14}$C-uric acid. Data shown are means ± SEM of at least three independent biological replicates (n=3-8). The statistical analysis was performed by unpaired two-tailed t-test. (*$P$< 0.05, **$P$< 0.01, ***$P$< 0.001, *vs.* WT). N.A., not active.

| **Mutations** | **$pIC_{50}$ (Mean ± SEM)** | | | |
|---|---|---|---|---|
| | **Benzbromarone** | **Dotinurad** | **Lesinurad** | **Verinurad** |
| WT | 6.71 ± 0.08 | 8.06 ± 0.06 | 4.64 ± 0.12 | 7.39 ± 0.07 |
| F241L | 7.28 ± 0.17 * | 7.03 ± 0.13 *** | 5.07 ± 0.13 * | 7.35 ± 0.13 |
| F360A | 5.74 ± 0.12 *** | 5.50 ± 0.36 *** | 4.79 ± 0.19 | 7.25 ± 0.13 |
| F364Y-F365Y | 5.14 ± 0.13 *** | 6.91 ± 0.18 *** | N.A. | 3.61 ± 1.32 ** |
| F449A | 5.44 ± 0.19 *** | / | 3.58 ± 0.25 ** | 6.90 ± 0.20 * |
| R477K | 6.38 ± 0.10 * | 7.64 ± 0.08 ** | 4.43 ± 0.09 * | 6.29 ± 0.07 *** |
